# Detecting Features of Protein Structure through their Mediator-Accessible Redox Activities

**DOI:** 10.1101/2023.12.12.571363

**Authors:** Dana Motabar, Eunkyoung Kim, Jinyang Li, Zhiling Zhao, Trina Mouchahoir, D. Travis Gallagher, John E. Schiel, Mamatha Garige, Carole Sourbier, Gregory F. Payne, William E. Bentley

**Affiliations:** Fischell Department of Bioengineering, University of Maryland, College Park, Maryland, United States; Institute for Bioscience and Biotechnology Research, Rockville, Maryland, United States; Robert E. Fischell Institute for Biomedical Devices, University of Maryland, College Park, Maryland, United States; National Institute of Standards and Technology, Gaithersburg, Maryland, United States; Laboratory of Molecular Oncology, Division of Biotechnology Review and Research-I, Office of Biotechnology Products, Office of Pharmaceutical Quality, Center for Drug Evaluation and Research (CDER), US Food and Drug Administration (FDA), Silver Spring, Maryland, United States

## Abstract

Protein function relies on sequence, folding, and post-translational modification, and molecular measurements are commonly used to reveal these structural features. Here, we report an alternative approach that represents these molecular features as readily measurable electronic patterns and validate this experimental approach by detecting structural perturbations commonly encountered during protein biomanufacturing. Specifically, we studied a monoclonal antibody standard (NISTmAb) and focused on the electronic detection of variants that have undergone interchain-disulfide bond reduction and methionine oxidation. Electronic detection of these structural perturbations is based on mediated electrochemical probing (MEP) that discerns patterns associated with the antibody’s mediator-accessible redox activity. We demonstrate that MEP can rapidly (within minutes) and quantitatively transduce the protein’s structural features into robust electronic signals that can enable bioprocess monitoring and control. More broadly, the ability to transduce information of a protein’s molecular structure into a more convenient electronic domain offers fundamentally new opportunities to apply the power of microelectronics and real-time data analytics to chemical and biological analysis.

## Introduction

The amino acid sequence of proteins encodes information regarding their structure, folding, and post-translational modifications. Commonly, the amino acids undergo redox (reduction-oxidation)-based modifications by redox active molecules (e.g., reactive oxygen species; ROS) which can impact protein structure and function. In some cases, these post-translational redox modifications may be essential to biological structure and function (e.g., thiol to disulfide sulfur switching of cysteine residues)^1–4^, while in other cases these modifications may be detrimental (e.g., oxidative damage)^5–7^.

Here, we report a rapid method to detect features of protein structure through their mediator-accessible redox activities (MARA), and we use therapeutic monoclonal antibodies (mAbs), as our model. Therapeutic antibodies are important based on their role in human health (therapeutics, diagnostics) and account for a significant portion of global sales revenue for all biopharmaceutical products^8–10^. Throughout their development and manufacture, mAbs can be altered by redox reactions resulting in interchain disulfide bond reduction or methionine oxidation which dramatically affect therapeutic efficacy, quality, and safety^11–13^.

As illustrated in **Fig. 1A**, antibody reduction is a common upstream bioprocessing issue that has been observed both in the bioreactor and during centrifugal harvest^14–16^. Reduction of the antibody’s disulfide bonds leads to lower molecular weight forms (containing free sulfhydryl groups) impacting product stability, bioactivity, and downstream process performance^13,17^. Irreversible oxidation of methionine residues can occur at all stages of bioprocessing and storage due to exposure to light, oxygen, or metal ions. Methionine oxidation also alters product quality and pharmacokinetic profiles^18,19^. Redox-based modifications are closely monitored to ensure optimal production, patient safety, and Food and Drug Administration (FDA) approval^20^.

**Fig. 1.**
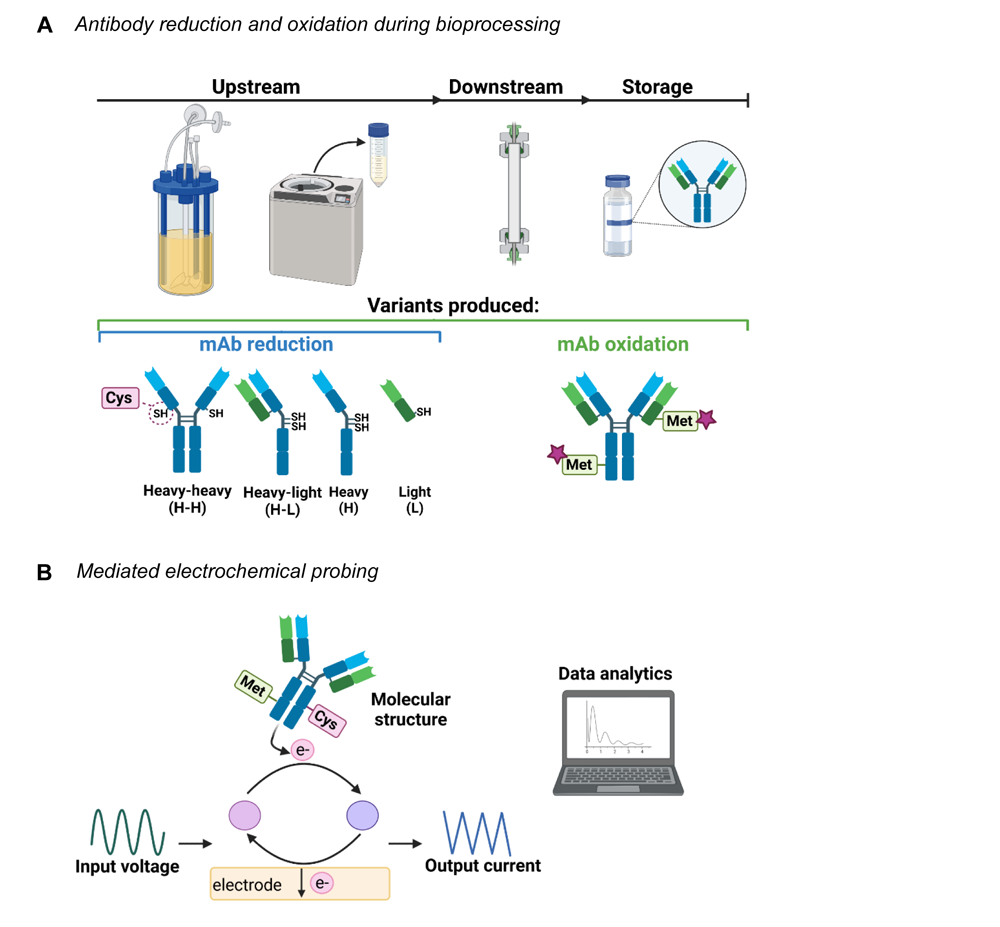
Mediated electrochemical probing to detect mAb variants. **(A)** During manufacturing and storage, redox reactions can yield mAb reduction and oxidation variants. **(B)** Mediated electrochemical probing measures the mAb’s mediator-accessible-redox-activity (MARA) and allows information of protein structure to be converted into a convenient electronic format.

Importantly, tools that enable access to redox based biological information are critical. “Gold standard” analytical methods (e.g., mass spectrometry, capillary electrophoresis) provide highly detailed chemical information which enables comprehensive and robust characterization of mAbs. However, these technologies tend to be expensive, labor intensive, and may require lengthy processing times. As such, it is well recognized that new process analytical methodologies are needed for therapeutic mAb production^21,22^, specifically those that allow for rapid, sensitive, and convenient measurement, and that can serve as a complement to more traditional analytical methods. These methodologies provide ready access to key process information and can be incorporated into advanced control schemes. We suggest that electrochemical measurements using electrodes are ideal for this purpose – the sterilizable pH and dissolved oxygen (DO) electrochemical probes are standard measurement tools in bioprocessing due to their reliability, ease of use, and ability to connect to electronics.

Here, we extend an emerging electrochemical method, mediated electrochemical probing (MEP) ^23–26^, for the simple, rapid, and sensitive detection of mAb reduction and oxidation variants. As illustrated in **Fig. 1B**, this method probes the mAb-containing sample using a diffusible redox mediator and an electrode with a precisely imposed voltage input sequence. In our case, the mediator is oxidized at the electrode and then diffuses into the sample where it probes the mAb for oxidizable residues. The MARA of the mAb is then detected when the reduced mediator returns to the electrode and its redox state is measured. Information of the mAb’s MARA is embedded in the electronic output features that are assessed using data analytics and comparative analysis^27,28^. Experimentally, we demonstrate the detection of mAb reduction and oxidation variants using a standard reference material (NISTmAb Reference Material 8671) and a commercial product, durvalumab. Overall, we believe this work demonstrates that mediated electrochemistry allows complex molecular information (e.g., of protein structure) to be rapidly converted into an electronic format that, in turn, enables broad application for on-site, remote, and deployable analysis.

## Results

### Reduction-probing the model amino acid, cysteine

To demonstrate MEP, we show how cysteine, one of the most important redox-sensitive molecular “switches” in biological systems, can be electrochemically characterized. When antibodies undergo reduction, the disulfide bonds are disrupted, and cysteine thiol groups are exposed. Our method leverages the exposed thiol groups to discern when the antibody is reduced. The scheme in **Fig. 2A** describes the proposed reaction mechanism for oxidative redox cycling between the redox mediator ferrocene dimethanol (Fc) and cysteine. In MEP, redox mediators (acting as electron shuttles) are added to biological samples which then interact with redox-active elements through the exchange of electrons^27,29^. When the appropriate input voltage is applied, Fc is oxidized by the electrode (Fc^Ox^), diffuses away from the electrode surface, and exchanges electrons with cysteine. This redox cycling reaction regenerates the reduced form (Fc^Red^) due to the transfer of electrons from cysteine resulting in an amplification in the oxidative current relative to the mediator alone in solution. Conversely, in the second panel of **Fig. 2A**, attenuation in the reductive current occurs due to the decrease in availability of Fc^Ox^.

**Fig. 2.**
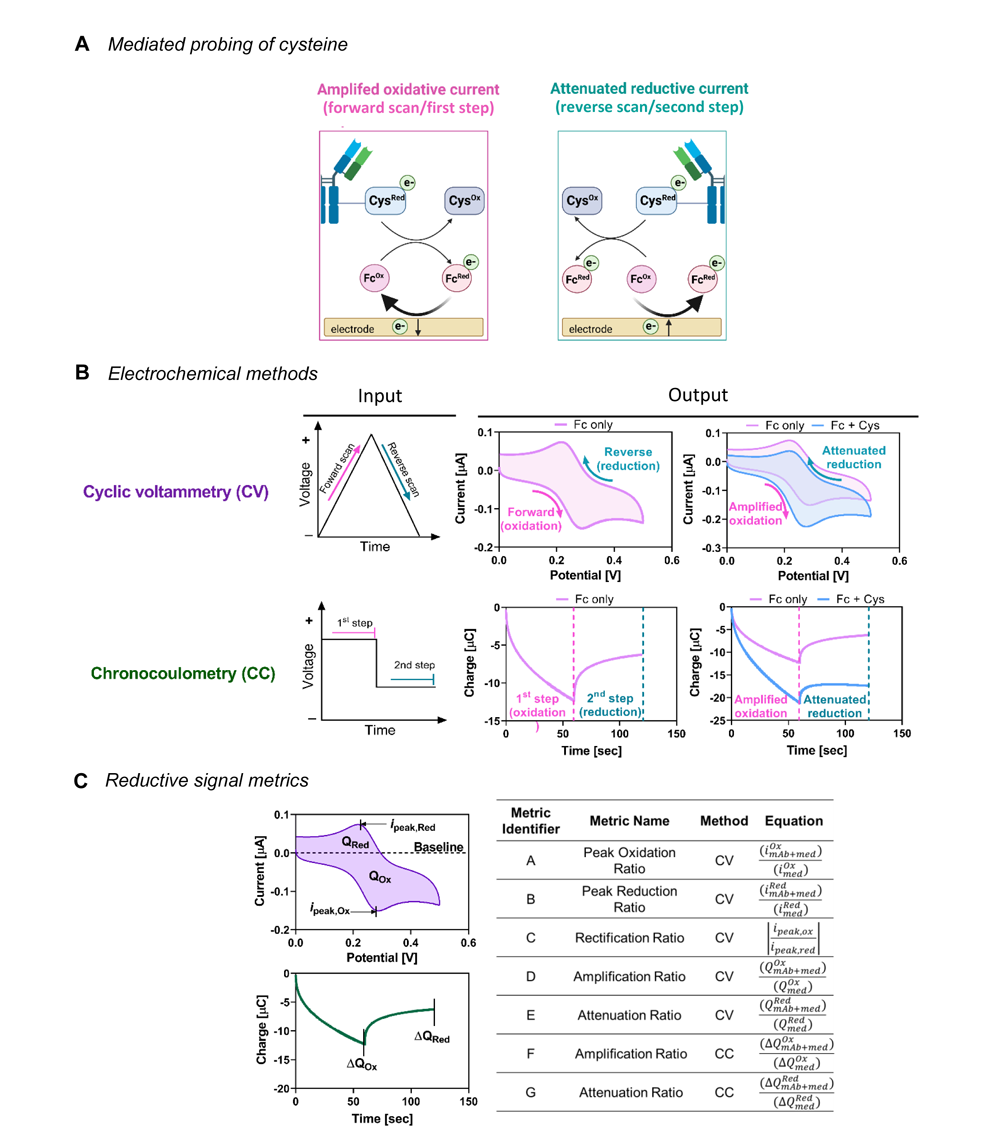
Mediated probing of cysteine. **(A)** Proposed reaction mechanism during Fc mediated redox cycling. **(B)** CV and CC plots of mediated probing of cysteine (50 µmol/L) with Fc (50 µmol/L). **(C)** Plots and table describe the reductive signal metrics for quantitative analysis.

As illustrated in **Fig. 2B**, Fc-mediated oxidation of cysteine can be measured using either of two electrochemical methods with distinct voltage input schemes, cyclic voltammetry (CV) or chronocoulometry (CC). For CV, as shown with the Fc (50 μmol/L) control, the input potential is scanned in the forward (positive) direction which leads to an oxidative peak (Fc^Red^ → Fc^Ox^ + e^−^) and then scanned in the reverse (negative) direction which leads to a reductive peak (Fc^Ox^ + e^−^ → Fc^Red^). For CC, the applied potential is fixed, but the principle is the same: quantification of the current generated by redox reactions occurring at the applied voltage. Here, the first voltage input was a 1 min step change to an oxidative value which oxidized Fc (+0.4 V vs. Ag/AgCl) and the second voltage input was a 1 min step change to a reductive value which reduced the oxidized Fc (+0.1 V vs. Ag/AgCl). When cysteine (50 μmol/L) was added to Fc (50 μmol/L), the CV and CC responses were dramatically altered as compared to the Fc controls. Specifically, the oxidative responses were significantly amplified, and the reductive responses were attenuated, following from Fig. 2A. When reduced cysteine is present, Fc oxidized at the electrode exchanges electrons with cysteine, becomes reduced, and then returns to the electrode to be oxidized. The current generated is the oxidative peak. The reaction with cysteine (reactive sink) enables greater current generated by Fc/electrode interactions than in the absence of cysteine (see controls).

Importantly, because both the CV and CC have characteristic shapes and are highly reproducible, signal metric analyses can be applied to quantitatively analyze the data. We suggest this is analogous to the assessment of cardiac function with electrocardiogram (EKG) measurements in which electrodes are used to discern a signature pattern that characterizes the health and functioning of the heart^30^. **Fig. 2C** lists the parameters and calculations by which our metrics are defined. For CV, the oxidative and reductive charges (Q_Ox_ or Q_Red_) were defined as the area under the curve below or above the zero-current baseline, respectively. The oxidative and reductive peak currents (*i_p_*_eak,ox_ or *i*_peak,red_) are the minima and maxima in the CV peak currents, respectively. For CC, the oxidative charge (ΔQ_Ox_) is defined as change in the charge after the 1^st^ minute and the reductive charge (ΔQ_Red_) is defined as the difference between the charges of the 1^st^ and 2^nd^ minutes. Metrics that are based on the amplification of the oxidative response were expected to increase (i.e., metrics A, C, D, and F) in the presence of reduced cysteine due to the free thiol content whereas metrics based on the attenuation of the reductive response were expected to decrease (i.e., metrics B, E, and G). For brevity, the metrics will later be referred to by their identifier (metrics A-G; see **Fig. 2C**).

### Detection of antibody reduction using mediated probing

Next, we investigated the ability of MEP to discern between intact and reduced NISTmAb. We first generated the reduced NISTmAb intermediate species using the reduction protocol illustrated in **Fig. S1A**. Tris(2-carboxyethyl)phosphine (TCEP) was added to NISTmAb (5 g/L) to a final TCEP concentration of 10 mmol/L, incubated for 4 h, then dialyzed into PBS for further analysis. Generally, Ellman’s assay and microchip capillary electrophoresis (CE) are used to characterize antibody reduction. Further details are provided in the Supplementary Information. In brief, Ellman’s assay (**Fig. S1B)** confirmed the presence of free thiol groups in the reduced samples across all concentrations evaluated (3 g/L, 1.5 g/L, 0.75 g/L and 0.25 g/L). Importantly, the increase in mAb concentration was directly correlated with increased free thiol content. Microchip CE (**Fig. S1C**) was also used to evaluate free heavy and light chain monomers and intermediate species generated upon antibody reduction. The CE results demonstrated that the reduction protocol was effective as there was minimal intact mAb and high levels of fully reduced heavy and light chains. We also note that such intermediate species appear during reduction events in upstream bioprocessing settings^15,17^. Importantly, these data confirm that the TCEP reduction protocol results in antibody fragments that contain free thiol groups.

For electrochemical analysis, Fc (50 μmol/L) was added to intact and reduced NISTmAb samples. As shown in the CV and CC plots of **Fig. S2A**, we first determined that there were no significant alterations in the electrochemical responses of the Fc mediator in the presence of intact mAb (Fc vs. Fc + intact). Furthermore, buffer controls for intact and reduced mAb (**Fig. S2B**) verified that the TCEP was effectively removed and that the background responses between the samples were similar. TCEP is known to have has its own electrochemical signature ^31^ and these data illustrate its absence after dialysis. The top panels of **Fig. 3A** show the results of MEP analysis of Fc (50 μmol/L) mixed with a range of industrially relevant mAb concentrations (3 g/L, 1.5 g/L, 0.75 g/L, and 0.25 g/L). As expected, we observed that the intact mAb responses (i.e., pink shades) remained essentially unchanged across all concentrations evaluated. In contrast, for the reduced mAb, we observed: (i) redox cycling between Fc and the reduced mAb as noted by increased oxidative current and decreased reductive current at the peak potentials and, (ii) higher oxidative amplification and reductive attenuation as the reduced mAb concentration increased. The concentration-dependent alterations in the CV and CC responses can be correlated with the free thiol content of the samples (see **Fig. S1B**, table).

**Fig. 3.**
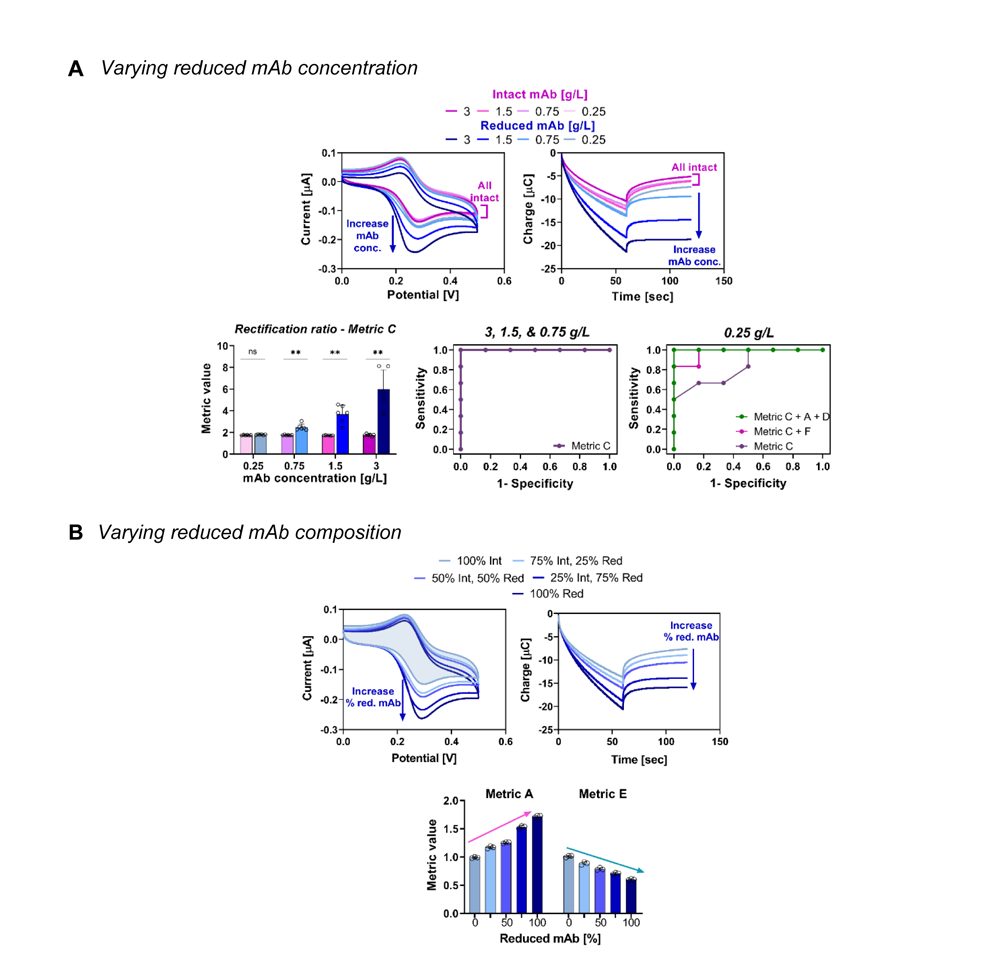
Varying reduced mAb concentration and composition. **(A)** Representative CV and CC of intact and reduced mAb at different concentrations: 3 g/L, 1.5 g/L, 0.75 g/L, and 0.25 g/L (*n* = 6). Corresponding signal metrics plot and ROC curves. Data was analyzed using the Mann-Whitney test (bar graph) and a logistic regression model (ROC plot). For bar graph, statistical significance defined as: * = *p* ≤ 0.05, ** = p ≤ 0.01, ns = *p* > 0.05. For ROC curve of 0.25 g/L NISTmAb, the discriminatory power of the model increases as additional metrics are combined: metric C (AUC = 0.833), metrics C + F (AUC = 0.972), and metrics C + A + D (AUC = 1.000). **(B)** Representative CV and CC plots of ratios of intact to reduced mAb (total mAb, 3 g/L; *n* = 3): 100% intact, 75% intact/25% reduced, 50% intact/50% reduced, 25% intact/75% reduced, and 100% reduced. Corresponding plot of best fit signal

We applied signal metric analyses to further confirm that observed differences between intact and reduced mAbs were statistically significant. First, the Mann-Whitney test was used to evaluate whether the calculated metrics for the two groups were distinct (*n* = 6 for both groups). In the bar graphs of **Fig. 3A**, metric C provided clear discriminatory power among all samples at higher concentrations (3 g/L, 1.5 g/L, and 0.75 g/L). Further, we found that all the calculated metrics enabled powerful discrimination, particularly at the higher concentrations (**Fig. S3A**). Further, **Fig. S3B** shows that simple linear regression analyses revealed concentration dependence (R^2^ ≥ 0.95) for all metrics across all concentrations. Interestingly, we also found no correlation with intact mAb for nearly all metrics (outliers, metrics E and G), which is consistent with the earlier result that Fc does not redox cycle with intact mAb, presumably because it has a full set of disulfide bonds.

Akin to evaluating cardiovascular function wherein several metrics of an EKG are combined for comprehensive understanding, we applied a logistic regression model ^32^ to evaluate combinations of metrics to increase further the probability of discriminating between intact and reduced forms. Akaike Information Criterion (AIC) values were used to assess the goodness of fit and Receiver Operating Characteristic (ROC) curves with Area Under Curve (AUC) and *p*-values were used to assess the discriminating power. In **Fig. 3A**, at higher mAb concentrations (3 g/L, 1.5 g/L, and 0.75 g/L), a single metric (metric C) was sufficient to discriminate between intact and reduced samples (AUC = 1). However, combining metrics increased the AUC and the statistical significance (*p*-value) of the analysis so that even the lowest mAb concentration (0.25 g/L) could be discriminated. That is, when three metrics were used cooperatively (metric C, A, and D), the discriminating power improved (AUC = 1). Complete analyses are shown in **Fig. S4**.

We next tested the hypothesis that MEP can distinguish between intact and reduced mAb in samples comprised of varied concentrations of reduced and intact mAb. To assess, Fc (50 μmol/L) was added to NISTmAb mixed in the following ratios: 100% intact, 75% intact/ 25% reduced, 50% intact/50% reduced, 25% intact/75% reduced, 100% reduced (total mAb for all samples, 3 g/L). Representative CV and CC plots are shown in **Fig. 3B**. As the percent composition of reduced mAb increased, the oxidative response was amplified and the reductive response was attenuated, as anticipated. Clearly linear trends were revealed for metrics A and E (additional metrics shown in **Fig. S5A**). Upon further examination (**Fig. S5B**), we found that MEP metrics could not only distinguish intact from reduced mAb, but that results were concentration dependent and virtually all metrics showed linearity across the full range of mAb concentration evaluated.

Importantly, it would expand the usefulness of our methodology if samples could be analyzed in conditions where interfering components may be present. In bioprocessing settings, mAbs are in cell culture media containing vitamins, amino acids, and other potentially redox active components. To evaluate the degree to which the matrix affects our assay, NISTmAb (in PBS) was diluted in a 1:1 ratio (total mAb, 2 g/L) with either PBS, fresh media (AMBIC Basal Medium 1.1), or conditioned media (spent Ex-Cell Advanced Fed-Batch Medium from CHO cell cultivations). CV and CC plots (**Fig. 4A**) suggest that MEP consistently discerned between intact and reduced mAbs in each process setting. Signal metrics analyses (**Fig. 4B**) further indicated that for all conditions, all oxidative response metrics were observed to increase and all reductive response metrics decreased, as anticipated. While the discriminatory power of MEP was shown to be independent of the solution background, it is important to note that sample conditions do impact the magnitude of the electrochemical responses. For example, the reduced mAb in conditioned media had slightly lower values for metrics A, D, and F compared to samples in buffer or fresh media. Overlayed CV plots of samples in different backgrounds (**Fig. S6**), show how redox active components contribute to the CV or CC spectra as “signature” patterns deviate mainly from the Fc spectrum and less so from the blank solutions (denoted “Background”). Not shown are data from many experiments of varied mediator concentration, electrode material, and analyte concentration used to determine the most effective conditions as discussed below.

**Fig. 4.**
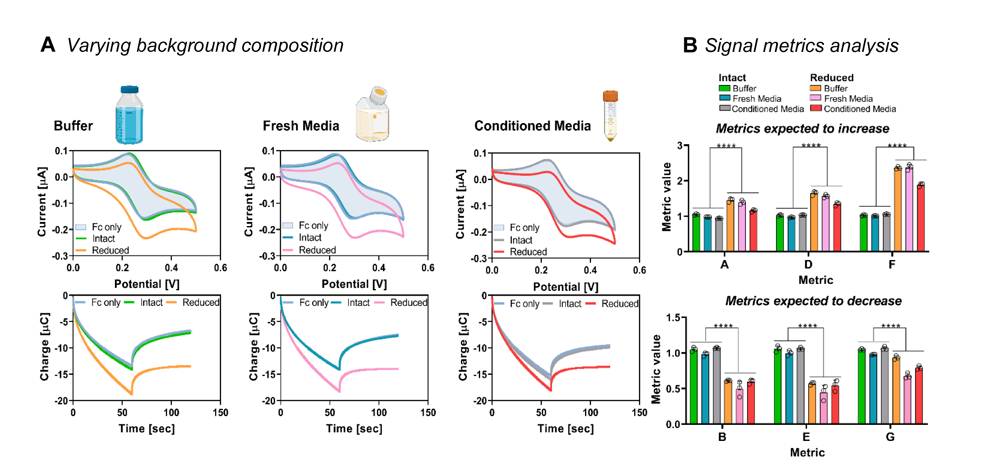
Reduced mAb in different background conditions. **(A)** Representative CV and CC plots of intact and reduced mAb which were diluted in a 1:1 ratio with phosphate buffered saline, fresh media, or conditioned media (total mAb, 2 g/L) (*n* = 3). **(B)** Signal metrics analysis shows that MEP can significantly discern intact from reduced mAb over all conditions tested. Data was analyzed using the Mann-Whitney test and statistical significance defined as **** = *p ≤* 0.0001. For all bar graphs, each bar represents the mean of the respective data set and error bars represent standard deviation.

In sum, by quantifying electrochemical features, calculating metrics, and statistically assembling their significance, MEP enables rapid and statistically valid assessment of mAb reduction.

### Oxidation - probing the model amino acid, methionine

Analogous to above targeting cysteine, we wanted to establish that MEP interacts with the primary target of antibody oxidation, methionine. Recognizing that methionine has a higher redox potential than cysteine, the redox mediator, hexachloroiridate(IV) (Ir), was used because it is a stronger oxidant than Fc (+0.67 V for Ir vs. +0.25 V for Fc). **Fig. 5A** describes the proposed reaction mechanism between Ir and methionine using MEP: when Ir (50 μmol/L) is added to methionine (2 mmol/L methionine; Ir + Met), redox cycling occurs resulting in significant amplification in the oxidative response for both CV and CC measurements (most notable at potentials above 0.6 V). When methionine is oxidized (2 mmol/L methionine sulfoxide added to buffer) and, therefore, electron deficient, redox cycling no longer occurs, and the oxidative response reverts to a level similar to that of the Ir control. In turn, Ir has no discernable reductive peak at these potentials. These data establish that Ir can redox cycle with methionine but not with oxidized methionine (no electron exchange). Hence, only oxidative signal metrics were applied to quantitatively analyze the results, potentially limiting discrimination. The CV and CC plots in **Fig. 5B** illustrate the parameters by which the metrics were defined. For CV, the oxidative charge (Q_Ox_) was defined as the area under the curve below the no current baseline and the peak oxidative current (*i*_peak,ox_) was defined as the maximum peak in the oxidative region. For CC, the potential was stepped to +0.85 V for 1-min, which oxidizes Ir and is defined as the oxidative charge (ΔQOx). The table (**Fig. 5B**, bottom) defines the equations and identifiers (1-6) of the metrics.

**Fig. 5.**
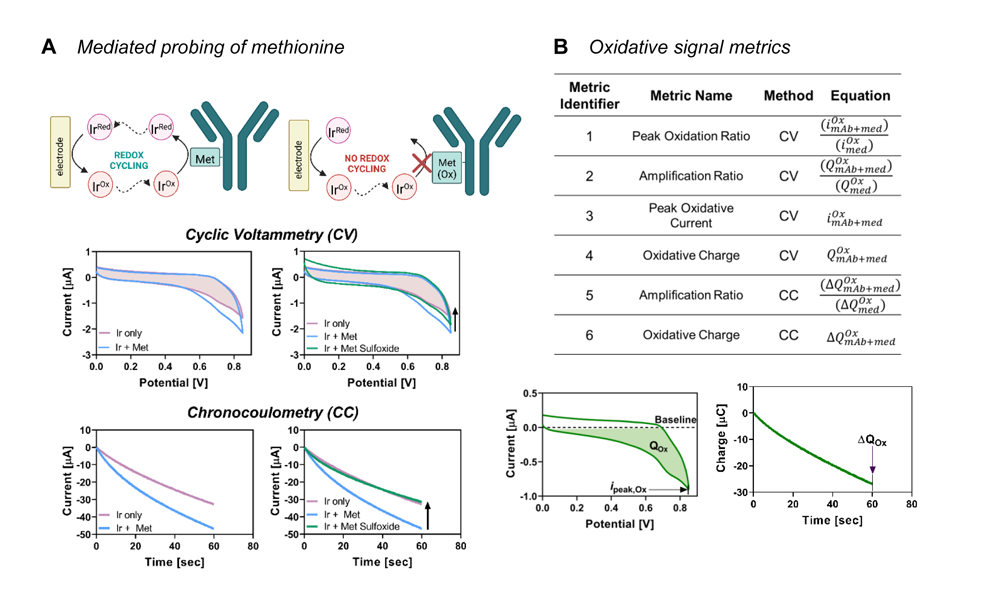
MEP of methionine. **(A)** Hexachloroiridate(IV) (Ir; 50 μmol/L) redox cycles with methionine (2 mmol/L). Ir does not cycle with the oxidized form of methionine (methionine sulfoxide; 2 mmol/L). **(B)** CV and CC plots along with corresponding table describe the oxidative signal metrics for quantitative analysis.

### Detection of antibody oxidation using mediated probing

We next investigated the ability of MEP to discern between control (non-oxidized) and oxidized mAbs. As before, we used NISTmAb, but here we also examined durvalumab, a clinically relevant immunotherapeutic. For oxidation of NISTmAb (**Fig. S7A**), hydrogen peroxide (1% v/v) was added to the mAb (4 g/L) and incubated in the dark for 48 h. This treatment preferentially oxidizes solvent exposed methionine residues^33–35^. Samples were then buffer exchanged into PBS prior to further analysis. Oxidation of the antibody’s methionine residues (identified for NISTmAb in **Fig. S7B**) was confirmed by LC-MS/MS peptide mapping. In brief, **Fig. S7C** reveals that six methionine residues (heavy chain M34, M101, M255, M361, M431; light chain M4) showed significant oxidation (> 95 %). The remaining two residues (heavy chain M87 and light chain M32) were only partially oxidized (< 7 %), likely due to their limited solvent accessibility (see Solvent Accessible Surface Area (SASA) analysis, **Fig. S7D**). Durvalumab was identically oxidized, and LC-MS/MS peptide mapping results (**Fig. S7E)** indicate that three of the five methionine residues (heavy chain M256, M362, M432) were significantly oxidized (> 97 %) while the two remaining residues (heavy chain M34 and M83) were only partially oxidized (< 3 %). Overall, these data confirm that the methionine residues of NISTmAb and durvalumab were oxidized due to incubation with hydrogen peroxide.

For electrochemical analysis, control and oxidized NISTmAb and durvalumab samples were diluted to 0.25 g/L, mixed with Ir (50 μmol/L), and assayed as above. In **Fig. S8A**, the oxidative response of the intact mAb (Ir + Intact) was amplified compared to the mediator by itself (Ir only). This demonstrates that Ir redox cycles with the surface exposed redox-active amino acids. The results from the CV and CC measurements for control and oxidized NISTmAb (**Fig. 6A**) reveal that oxidized mAb exhibited a decreased oxidative response compared to control mAb. While less dramatic, these data are consistent with the methionine results.

**Fig. 6.**
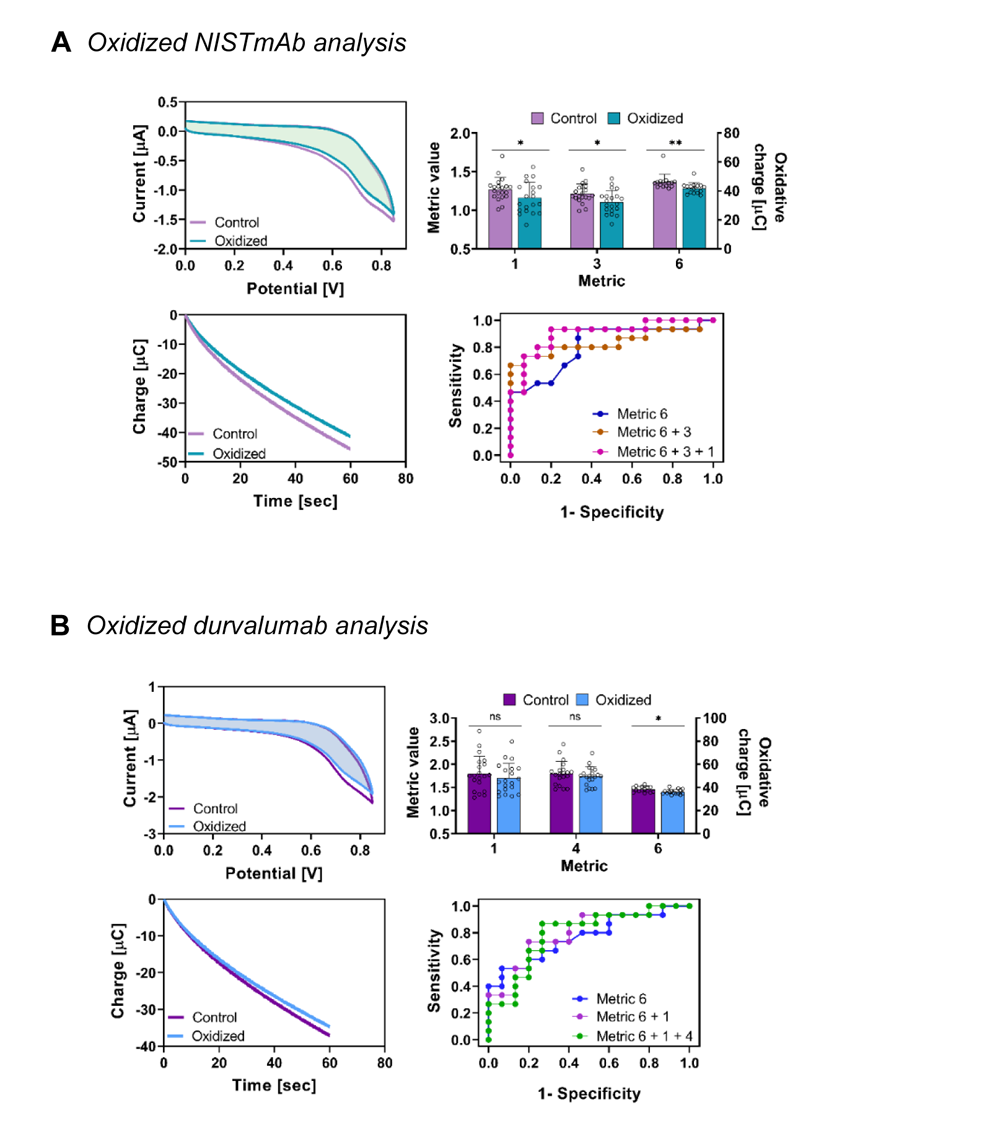
MEP of oxidized mAbs. **(A)** MEP of control and oxidized NISTmAb (0.25 g/L). For ROC plot, the discriminatory power of the model increases as additional metrics are combined: metric 6 (AUC = 0.813), metrics 6 + 3 (AUC = 0.840), and metrics 6 + 3 + 1 (AUC = 0.902). **(B)** MEP of control and oxidized durvalumab (0.25 g/L). Both mAbs are probed with Ir (50 μmol/L) using CV (*n* = 20) and CC (*n* = 15) techniques. For ROC plot, the combination of 2 metrics provided the optimal discriminatory power: metric 6 (AUC = 0.771), metrics 6 + 1 (AUC = 0.796), and metrics 6 +1 + 4 (AUC = 0.791). For bar charts, statistical significance defined as: * = *p* ≤ 0.05, ** = p ≤ 0.01, ns = *p* > 0.05. Data was analyzed using the Mann-Whitney test (bar graph) and a logistic regression model (ROC plot). For all bar graphs, each bar represents the mean of the respective data set and error bars represent standard deviation.

Mann-Whitney tests and logistic regression models were applied to evaluate the characteristic metrics. For the Mann-Whitney test, shown in the bar graphs of **Fig. 6A** and **Fig. S8B**, all metrics exhibited significantly different responses (*p =* * or **). We note, however, that a higher number of replicates was required to achieve these differences (CV: *n* = 20 and CC: *n* = 15 vs. *n* = 6 for mAb reduction studies). The logistic model again indicated that combinations of metrics increased resolving power (see ROC plot in **Fig. 6A** and the table in **Fig. S8C**). Here, a combination of multiple metrics (1, 3, and 6) increased the discriminating power of MEP for evaluating mAb oxidation.

As noted above, the mediator and mAb concentrations are important for effective MEP analysis. In **Fig. S9**, we varied the NISTmAb concentration (62.5 mg/L, 125 mg/L, 250 mg/L, 500 mg/L) while maintaining the same Ir concentration (50 μmol/L). Conversely, we also maintained the NISTmAb concentration (250 mg/L) while varying the Ir concentration (25 μmol/L, 50 μmol/L, 75 μmol/L, 100 μmol/L). We then quantitatively evaluated our data by applying select metrics (metrics 1 and 2, which had appeared most distinguishing). In both cases, we found that the same conditions (125 & 250 mg/L NISTmAb and 50 µmol/L Ir) resulted in the largest difference between control and oxidized samples. That is, at several other NISTmAb: Ir concentrations and ratios, we were unable to differentiate control from oxidized samples.

For durvalumab, which has five methionine residues (**Fig. S10A**), and using the identical concentrations as NISTmAb, we observed that the oxidative response of the control mAb (Ir + Control) was also amplified compared to the mediator by itself (Ir only). Results for metrics 1, 4 and 6 are shown in **Fig. 6B**. Like the NISTmAb, oxidized durvalumab had a lower average oxidative response compared to the intact protein in both CV and CC measurements. The statistical analysis, shown in bar graphs of **Fig. 6B** and **Fig. S10B**, with the Mann-Whitney test (using the same number of CV and CC replicates as NISTmAb) revealed that although the metrics trended as expected, only metric 6 had a statistically significant difference (*p =* *) between the groups. Interestingly, metric 6 also yielded the highest significant difference (*p =* **) for NISTmAb. For the logistic model, as displayed in the ROC plot and table in **Fig. S10C**, the combination of two metrics (metrics 6 + 1) improved the ability to distinguish between control and oxidized durvalumab (AUC = 0.796), while the addition of a third metric (metrics 6 + 1 + 4) had minimal impact (AUC = 0.791).

## Discussion

A protein’s functional attributes (e.g., activity and immunogenicity) are intimately linked to structure and thus measurements are integral to protein characterization. The challenge however is that protein structure extends over various length scales (from sequence to quaternary structure), and thus structural characterization often requires multiple methods of analysis (e.g., mass spectrometry and X-Ray crystallography). While these “gold standard” methods are essential for establishing structure-function relationships, they are slow and require specialized skills, making them difficult to adapt for near-real-time information that is often essential for decision-making in a biomanufacturing environment. Here, we report the use of mediated electrochemical probing (MEP) as an alternative – it rapidly converts a protein’s structural information directly into an electronic format. Specifically, MEP uses mediators and a precisely controlled potential (i.e., voltage) inputs to probe a protein’s mediator-accessible redox activity (MARA) and generates output response signals (i.e., currents) that can be correlated to relevant structural features.

First, we used the standard reference protein, NISTmAb, and examined how MEP could be used to detect partially reduced variants generated by interchain disulfide bond reduction. We observed that the weakly-oxidative Fc redox-mediator can selectively detect the free thiols of cysteine residues and that when variants with higher thiol contents were probed, **Fig. 3A and B** showed stronger MEP-based current responses. Importantly, the reduction of mAb disulfides that yields free thiols also results in partial dissociation of protein subunit chains. Thus, the MEP-based detection of free thiols in the reduced variants is also a measure of the loss of the mAb’s native quaternary structure. Second, we observed that MEP was able to discern between intact and reduced samples in the presence of interfering analytes (e.g., buffer, fresh media, conditioned media). As shown in **Fig. 4**, signal metrics trended similarly for samples in all three conditions. This is a major advantage of the MEP methodology as it can lower the processing time and effort required to measure reduced mAb, especially by lowering the number of time-consuming steps required for sample preparation.

Similarly, we used MEP to probe partially oxidized antibody variants using both the NISTmAb and durvalumab. In this case, we used a more strongly oxidative mediator, Ir. First, we showed that MEP detected statistically significant differences between the partially oxidized (vs. unoxidized) mAbs, and these differences appear correlated to solvent-exposed methionine residues that are common sites of mAb oxidation. **Fig. 6A and B** indicate that the lower significance in the differences between the oxidized and unoxidized mAb for durvalumab (vs. NISTmAb) may be related to the smaller number of methionine residues (three for durvalumab and six for NISTmAb). Importantly, these results suggest that the MEP methodology is sensitive to the amino acid composition of proteins. Second, we observed that detection of mAb oxidation variants was more challenging (than detecting reduction variants) and required greater experimental optimization and more extensive logistic regression analysis to increase the discerning power of the MEP-generated signals. Perhaps this is because of the number of MARA cysteine versus methionine residues in intact mAbs.

Overall, this work demonstrates the potential of MEP to convert information of molecular structure into a convenient electronic format. There are two important practical implications of this work. First, MEP measurements are made using simple, inexpensive and miniaturizable electrodes that could be deployed in manufacturing settings. Our specific example is relevant to the biomanufacturing of mAbs where we envision MEP could enable a new, device-based process analytical technology (PAT)^21^. Second, MEP provides near-real-time electronic data that allows instantaneous statistical analysis and decision-making. Thus, we envision MEP can provide timely, actionable-information of molecular-structure to enable process control in a biomanufacturing setting.

## Methods

### Materials

NISTmAb RM8671 (Lot 14HB-D-002, humanized IgG1κ) was obtained from National Institute of Standards of Technology (NIST) (Gaithersburg, MD, USA). Durvalumab (MedImmune/AstraZeneca) was purchased via McKesson Specialty Health (Scottsdale, Arizona). 1,1′-Ferrocenedimethanol (Fc) was purchased from Santa Cruz Biotechnology (Dallas, TX). Trizma HCl and Trizma base (collectively, Tris), guanidine HCl, ethylenediaminetetraacetic acid (EDTA), iodoacetamide (IAM), urea, recombinant porcine trypsin, K_2_IrCl_6_, and phosphate buffered saline (PBS, pH 7.4) were purchased from Sigma-Aldrich (St. Louis, MO). L-histidine monohydrochloride monohydrate and L-histidine were purchased from Avantor (Allentown, PA). 0.1 % formic acid was purchased from Honeywell (Phoenix, AZ). 5 mmol/L stock solutions of Fc and K_2_IrCl_6_, respectively, were prepared in PBS and aliquots were stored at −20 °C. TCEP (tris(2-carboxyethyl)phosphine) bond breaker solution, Zeba Spin Desalting columns (7KDa MWCO), Slide-A-Lyzer Dialysis Cassettes (10K MWCO), 5,5-dithio-bis-(2-nitrobenzoic acid) (DTNB, aka Ellman’s Reagent) and DL-dithiothreitol (DTT) were purchased from Thermo Fisher Scientific (Waltham, MA). AMBIC Basal Medium 1.1 (Lonza, Basel, Switzerland) was provided by the Advanced Mammalian Biomanufacturing Innovation Center (AMBIC).

### Antibody reduction protocol

NISTmAb samples (10 g/L stock) were first diluted to 5 g/L with PBS, pH 7.4. For reduced samples, TCEP was added to 1 mL of NISTmAb for a final TCEP concentration of 10 mmol/L. The intact and reduced samples were then incubated for 4 hours at room temperature. Both intact and reduced samples were dialyzed using Slide-A-Lyzer dialysis cassettes (0.5 mL to 3 mL, 10KDa MWCO) in 2 L of PBS, pH 7.4 overnight at 4 °C. After dialysis, samples were stored at −20 °C until analysis. Buffer controls were performed using 25 mmol/L histidine buffer, pH 6.0 with and without the addition of TCEP. The controls were diluted with PBS, pH 7.4 in an identical manner to experimental samples.

### Determination of antibody reduction by standard approaches

To analyze antibody reduction, microchip capillary electrophoresis (CE) was performed using a 2100 Bioanalyzer (Agilent). Intact and reduced NISTmAb samples were run using the Agilent Protein 230 kits under non-reducing conditions according to the manufacturer’s protocols. A sulfhydryl assay was performed to determine free thiol content in reduced antibody samples. Briefly, 625 μL of reaction buffer (0.1 mol/L sodium phosphate, 1mmol/L EDTA, pH 8) was mixed with 12.5 μL of 4 mg/mL of DTNB solution (in reaction buffer). 62.5 μL of intact or reduced NISTmAb (3 g/L, 1.5 g/L, 0.75 g/L, and 0.25 g/L, dilutions made with PBS) were added to the mixture and samples were then incubated for 15 min at room temperature. Spectral absorption at 412 nm was measured and the free thiol content was calculated using the molar extinction coefficient of TNB (14,150 L mol^-1^ cm-^1^) according to the manufacturer’s instructions.

### Antibody oxidation protocol

NISTmAb stock (10 g/L) and durvalumab stock (50 g/L) were first diluted to 4 g/L in PBS. To oxidize the antibodies, hydrogen peroxide (1% v/v) was added. Both control and oxidized samples were incubated for 48 h in the dark at room temperature. The samples were then buffer exchanged into PBS using Zeba Spin Desalting columns (7KDa MWCO). Samples were then stored at −20 °C until analysis. Buffer controls were performed using 25 mmol/L histidine buffer, pH 6.0 with and without the addition of hydrogen peroxide. The controls were diluted with PBS, pH 7.4 in an identical manner to experimental samples.

### LC-MS/MS peptide mapping

Control and oxidized sample concentrations were diluted to 1.9 µg/µL with denaturing buffer (5.4 mol/L guanidine HCl, 0.9 mmol/L EDTA, 90 mmol/L Tris, pH 7.8). Samples were reduced by adding DTT to a final concentration of 15 mmol/L, followed by incubation at 4 °C for 1 h. Samples were then alkylated by adding IAM to a final concentration of 29 mmol/L, followed by incubation at 4 °C for 1 h in the dark. Samples were exchanged to digestion buffer (1 mol/L urea in 0.13 mol/L Tris) using Zeba spin columns per the manufacturer’s instructions. Trypsin was added to the samples for a final enzyme: IgG mass ratio of 1:18, followed by incubation at room temperature for 4 h. Finally, 0.1 % formic acid was added at a 1:1 volume ratio to each sample. Digested samples were stored at – 80 °C until analysis. 5 µg of peptide digests were loaded via autosampler onto a C18 column and analyzed by liquid chromatography (LC)/ mass spectrometry (MS). Peptide identification and quantification of methionine oxidation were performed using Genedata Expressionist v 15.0.6.

### Solvent-accessible surface area analysis

Solvent-exposed areas of sulfurs in methionine residues in structures 5K8A (the NISTmAb Fab, with 5 Mets) and 5VGP (the NISTmAb Fc, with 3 Mets) were calculated by AREAIMOL in the CCP4 software suite^36^, neglecting protein hydrogens and using default van der Waal radii. To account for flexibility, sulfur exposure numbers were averaged with their adjacent carbons, and weighted for sequence proximity to peptide termini (only one Met was near enough to affect its exposure value). The resulting structure-based estimates of peroxide exposure have correlation 0.87 with oxidation measurements (Fig. S7D).

### Determination of antibody reduction by mediated electrochemical probing

Cyclic voltammetry (CV) and chronocoulometry (CC) measurements were taken using a CHI1040C electrochemical analyzer (CH Instruments; Austin, TX). The measurements were performed with a 3-electrode system containing a 2 mm diameter glassy carbon working electrode, a Pt wire serving as the counter electrode, and an Ag/AgCl reference electrode. For electrochemical measurement, samples were diluted to the appropriate concentration and then Fc (50 μmol/L) was added. For fresh media studies, NISTmAb (total mAb, 2 g/L) was spiked into AMBIC 1.1 Basal Medium. For conditioned media studies, NISTmAb (total mAb, 2 g/L) was spiked into spent Ex-Cell Advanced Fed-Batch Medium media (cells removed) from CHOZN23 (MilliporeSigma) cell culture. CV measurements were taken over a potential range of 0 V to 0.5 V at a scan rate of 2 mV/s. Chronocoulometry measurements were taken at a constant potential of +0.4 V for 1 min and then switched to a reductive potential of +0.1 V for 1 min. All electrochemical measurements were taken from distinct samples. The working electrode was polished using 0.05 μm alumina and rinsed with ddH_2_O between each measurement. All potentials are reported versus Ag/AgCl.

### Determination of antibody oxidation by mediated electrochemical probing

The electrochemical measurements were performed the same as above but with the following modifications. NISTmAb and durvalumab samples were first diluted to the appropriate concentrations and then potassium hexachloroiridate(IV) (50 μmol/L) was added to samples. CV measurements were taken over a potential range of 0 V to 0.85 V at a scan rate of 10 mV/s. CC measurements were taken at a constant input potential of +0.85 V for 1 min. All electrochemical measurements were taken from distinct samples. The working electrode was polished using 0.05 μm alumina and rinsed with ddH_2_O between each measurement. All potentials are reported versus Ag/AgCl.

### Statistical analysis

Statistical analysis was performed with Prism Graphpad and R version 3.4.3. Differences between intact and reduced or oxidized antibodies were assessed using the Mann-Whitney U test (two-sided). A logistic regression model (‘nnet’ package) was used to assess the combination effect of multiple signal metrics on differentiating intact from reduced or control from oxidized mAbs. The Akaike Information Criterion (AIC) was used to estimate the quality of each model. The discriminating ability was evaluated using ROC (Receiver Operating Characteristic) metric with AUC ((Area Under Curve) (i.e., c-statistic)) and *p*-values.

## Acknowledgements

This work was supported by the Defense Threat Reduction Agency (HDTRA1-19-1-0021), the Advanced Mammalian Biomanufacturing Innovation Center (IUCRC2100632, #2004614245), and the National Science Foundation (MCB #2227598). This work was also supported in part by a Center of Excellence in Regulatory Science and Innovation (CERSI) grant to University of Maryland from the US Food & Drug Administration (#5U01FD005946-04). This article reflects the views of the authors and should not be construed to represent U.S. FDA’s views or policies. The authors declare no conflict of interest. Identification of commercial materials and equipment does not imply recommendation or endorsement by the National Institute of Standards and Technology. Figures were created in part with BioRender.com.

## Data availability statement

The authors declare that the data supporting the findings of this study are available within the paper and its Supplementary Information file. If raw data files are needed in another format, they are available from the corresponding author upon reasonable request.

## Author contributions

Methodology: DM, EK, JL, ZZ, TM, TDG, JES, MG, CS, GFP, WEB

Visualization: DM, EK, TM, TDG

Writing, review & editing: DM, EK, TM, TDG, MG, CS, GFP, WEB

