## Supplemental File for "Detecting Features of Protein Structure through their Mediator-Accessible Redox Activities"

**A** *Reduction protocol*

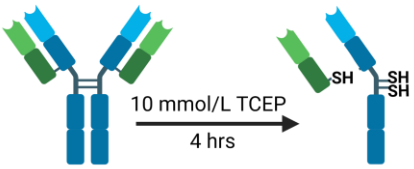


**B** *Ellman's assay for reduction*

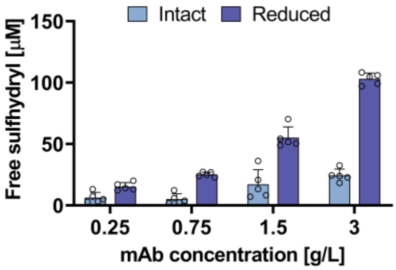


| Concentration [g/L] | Ellman's Assay free sulfhydryl [μM] Intact | Ellman's Assay free sulfhydryl [μM] Reduced | *Theoretical* free sulfhydryl [μM] Reduced |
|---|---|---|---|
| 0.25 | 6.4 ± 4.2 | 15.3 ± 3.2 | 9.4 |
| 0.75 | 5.2 ± 4.2 | 25.2 ± 1.9 | 28.1 |
| 1.5 | 17.1 ± 12.0 | 55.2 ± 8.7 | 56.2 |
| 3.0 | 24.4 ± 5.4 | 103.1 ± 4.7 | 112.3 |

**C** *CE analysis for reduction*

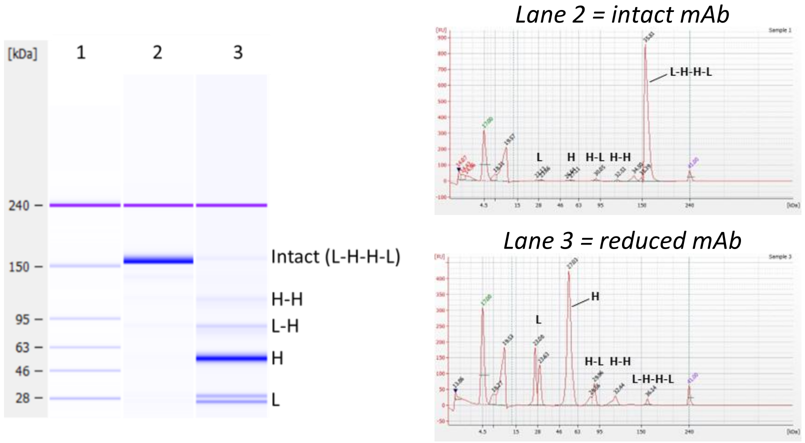


| Lane 3 species | MW [kDa] | Intact [%] | Reduced [%] |
|---|---|---|---|
| Intact (L-H-H-L) | 150 | 94.7 | 1.3 |
| HH | 100 | 1.3 | 4.3 |
| HL | 75 | 1.3 | 8.9 |
| H | ~50 | 0.8 | 57.0 |
| L | ~25 | 1.9 | 28.5 |

**Fig. S1. Methods to create and analyze reduced mAb variants. (A)** Method used for antibody reduction. **(B)** Ellman's assay results confirm that the reduced mAb at all concentrations evaluated (3 g/L, 1.5 g/L, 0.75 g./L, and 0.25 g/L) contain elevated free thiol concentrations as compared to the corresponding intact samples ($n$ = 5). Error bars in plot represent the standard deviation. **(C)** Simulated gel and corresponding electropherograms from microchip capillary electrophoresis show the types of mAb species produced from reduction. For gel: lane 1 = ladder, lane 2 = intact mAb, lane 3 = reduced mAb. As expected, a single, bold band (lane 2) around ~150 kDa represents the intact (non-reduced) antibody. Conversely, for the reduced antibody (lane 3), there is a faint band representing the remaining intact antibody (~150 kDa) and several lower molecular weight reduction species representing light chain (L; ~25 kDa), heavy chain (H; ~50 kDa), L-H intermediate (75 kDa), and H-H intermediate (100 kDa). After the reduction protocol, there was minimal intact mAb (1.3%) and high levels of reduced species, with free heavy chain comprising the greatest amount (57%).

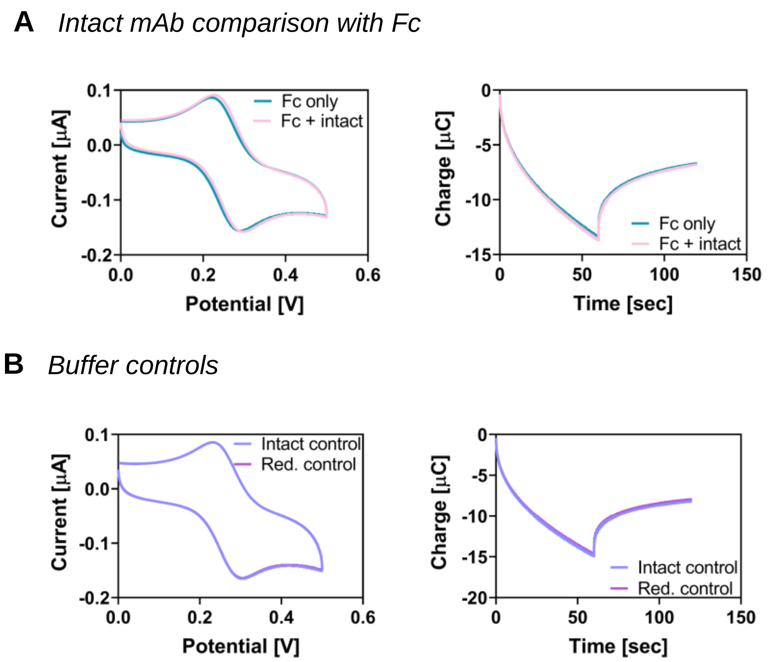


**Fig. S2. Controls for reduced mAb. (A)** CV and CC plots show that there is minimal difference between Fc only (50 µM) and Fc with intact mAb (3 g/L). **(B)** CV and CC plots of buffer controls for intact and reduced mAb. Intact buffer control was 25 mmol/L Histidine-HCL, pH 6.0 and reduced buffer control was 25 mmol/L Histidine-HCL,10 mmol/L TCEP, pH 6.0. In an identical manner to the mAb-containing samples, buffer controls were dialyzed overnight using Slide-A-Lyzer dialysis cassettes into phosphate buffered saline, pH 7.4. CV and CC measurements were taken which showed that there is no significant difference between intact and reduced buffer controls.

**A** *Signal metrics analysis*

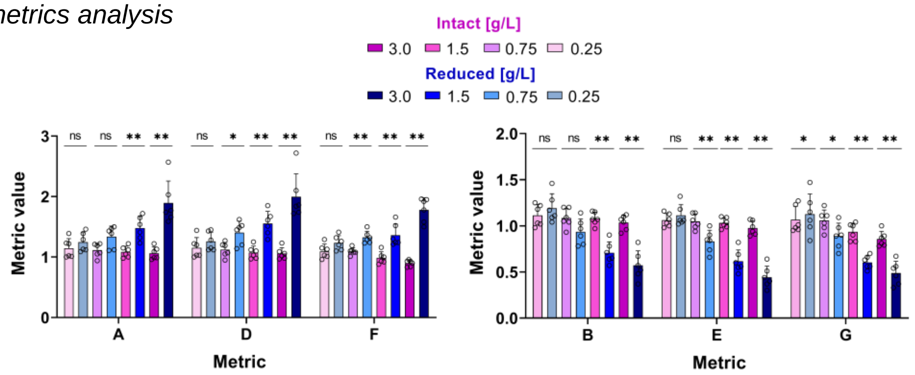


### *3 g/L*

| Metric | Intact | Reduced | *p* -value |
|---|---|---|---|
| A | 1.06 ± 0.10 | 1.89 ± 0.36 | ** |
| B | 1.03 ± 0.08 | 0.57 ± 0.16 | ** |
| C | 1.76 ± 0.10 | 6.01 ± 1.77 | ** |
| D | 1.06 ± 0.09 | 2.00 ± 0.38 | ** |
| E | 0.98 ± 0.08 | 0.44 ± 0.12 | ** |
| F | 0.90 ± 0.05 | 1.74 ± 0.27 | ** |
| G | 0.86 ± 0.08 | 0.39 ± 0.23 | ** |

### *1.5 g/L*

| Metric | Intact | Reduced | *p* -value |
|---|---|---|---|
| A | 1.09 ± 0.09 | 1.48 ± 0.19 | ** |
| B | 1.08 ± 0.0 | 0.71 ± 0.12 | ** |
| C | 1.72 ± 0.02 | 3.68 ± 0.84 | ** |
| D | 1.08 ± 0.10 | 1.55 ± 0.20 | ** |
| E | 1.03 ± 0.05 | 0.62 ± 0.12 | ** |
| F | 0.98 ± 0.10 | 1.36 ± 0.18 | ** |
| G | 0.93 ± 0.09 | 0.60 ± 0.07 | ** |

### *0.75 g/L*

| Metric | Intact | Reduced | *p* -value |
|---|---|---|---|
| A | 1.11 ± 0.11 | 1.34 ± 0.18 | ns |
| B | 1.09 ± 0.10 | 0.93 ± 0.14 | ns |
| C | 1.76 ± 0.03 | 2.47 ± 0.30 | ** |
| D | 1.12 ± 0.12 | 1.40 ± 0.19 | * |
| E | 1.05 ± 0.08 | 0.83 ± 0.12 | ** |
| F | 1.10 ± 0.06 | 1.30 ± 0.10 | ** |
| G | 1.06 ± 0.11 | 0.89 ± 0.14 | * |

### *0.25 g/L*

| Metric | Intact | Reduced | *p* -value |
|---|---|---|---|
| A | 1.15 ± 0.16 | 1.25 ± 0.16 | ns |
| B | 1.11 ± 0.12 | 1.19 ± 0.15 | ns |
| C | 1.76 ± 0.02 | 1.80 ± 0.04 | ns |
| D | 1.15 ± 0.17 | 1.26 ± 0.15 | ns |
| E | 1.06 ± 0.07 | 1.11 ± 0.12 | ns |
| F | 1.09 ± 0.12 | 1.24 ± 0.12 | ns |
| G | 1.07 ± 0.15 | 1.13 ± 0.22 | * |

**B** *Linear regression analysis*

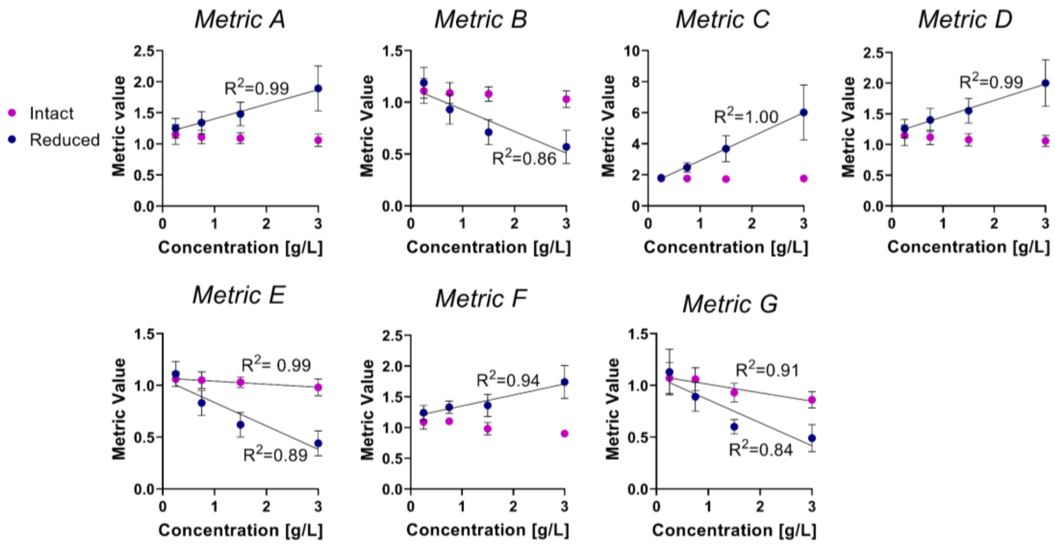


**Fig. S3. Varying reduced mAb concentration**. **(A)** Bar graphs and tables of signal metrics derived from corresponding CC and CV data in Figure 3A. **(B)** Linear regression of CV and CC signal metrics derived from Figure 3A CC and CV data across all reduced mAb concentrations evaluated. For intact samples (pink) the absence of a regressed line indicates slope was not significantly different than zero. For Metric E, the upper bound of the < $p$ = 0.05 confidence limit was 0.007. This slope and that Metric G, were deemed significantly different than zero. Error bars in all plots represent the standard deviation.

### *3 g/L*

| # of metrics | Combination | AIC | AUC | *p*-value |
|---|---|---|---|---|
| 1 | Metric A | 4.06 | 1.00 | 0.0050 |
| | Metric B | 4.26 | 1.00 | 0.0043 |
| | **Metric C** | **4.00** | **1.00** | **0.0037** |
| | Metric D | 4.03 | 1.00 | 0.0022 |
| | Metric E | 4.04 | 1.00 | 0.0050 |
| | Metric F | 4.02 | 1.00 | 0.0047 |
| | Metric G | 4.35 | 1.00 | 0.0050 |
| 2 | Metric C+ A | 6.00 | 1.00 | 0.0037 |
| | Metric C+ B | 6.00 | 1.00 | 0.0034 |
| | Metric C+ D | 6.00 | 1.00 | 0.0037 |
| | Metric C+ E | 6.00 | 1.00 | 0.0037 |
| | Metric C+ F | 6.00 | 1.00 | 0.0037 |
| | Metric C+ G | 6.00 | 1.00 | 0.0037 |
| | Metric F + A | 6.00 | 1.00 | 0.0043 |
| | Metric F + B | 6.00 | 1.00 | 0.0050 |
| | Metric F + C | 6.00 | 1.00 | 0.0037 |
| | Metric F + D | 6.00 | 1.00 | 0.0037 |
| | Metric F + E | 6.00 | 1.00 | 0.0050 |
| | Metric F + G | 6.00 | 1.00 | 0.0022 |
| 3 | Metric F + G + C | 8.00 | 1.00 | 0.0037 |
| | Metric F + G + D | 8.00 | 1.00 | 0.0050 |
| 4 | Metric C + F + E + D | 10.00 | 1.00 | 0.0037 |
| 7 | Metric A + B + C + D + E + F + G | 16.00 | 1.00 | 0.0043 |

### *1.5 g/L*

| # of metrics | Combination | AIC | AUC | *p*-value |
|---|---|---|---|---|
| 1 | Metric A | 6.42 | 1.00 | 0.0050 |
| | Metric B | 5.01 | 1.00 | 0.0050 |
| | **Metric C** | **4.05** | **1.00** | **0.0041** |
| | Metric D | 5.75 | 1.00 | 0.0022 |
| | Metric E | 4.15 | 1.00 | 0.0048 |
| | Metric F | 5.25 | 1.00 | 0.0050 |
| | Metric G | 4.23 | 1.00 | 0.0050 |
| 2 | Metric C + A | 6.00 | 1.00 | 0.0037 |
| | Metric C + B | 6.00 | 1.00 | 0.0037 |
| | Metric C + D | 6.00 | 1.00 | 0.0037 |
| | Metric C + E | 6.00 | 1.00 | 0.0037 |
| | Metric C + F | 6.00 | 1.00 | 0.0037 |
| | Metric C + G | 6.00 | 1.00 | 0.0037 |
| 7 | Metric A + B + C + D + E + F + G | 16.00 | 1.00 | 0.0023 |

### *0.75 g/L*

| # of metrics | Combination | AIC | AUC | *p*-value |
|---|---|---|---|---|
| 1 | Metric A | 14.87 | 0.833 | 0.0650 |
| | Metric B | 16.41 | 0.833 | 0.0650 |
| | **Metric C** | **4.03** | **1.00** | **0.0650** |
| | Metric D | 14.87 | 0.833 | 0.0650 |
| | Metric E | 16.41 | 0.833 | 0.0650 |
| | Metric F | 5.17 | 1.00 | 0.0050 |
| | Metric G | 15.68 | 0.861 | 0.0440 |
| 2 | Metric C + A | 6 | 1.00 | 0.0050 |
| | Metric C + B | 6 | 1.00 | 0.0043 |
| | Metric C + D | 6 | 1.00 | 0.0050 |
| | Metric C + E | 6 | 1.00 | 0.0043 |
| | Metric C + F | 6 | 1.00 | 0.0050 |
| | Metric C + G | 6 | 1.00 | 0.0036 |
| 3 | Metric C + F + G | 8 | 1.00 | 0.0034 |
| 7 | Metric A + B + C + D + E + F + G | 16 | 1.00 | 0.0034 |

### *0.25 g/L*

| # of metrics | Combination | AIC | AUC | *p*-value |
|---|---|---|---|---|
| 1 | Metric A | 19.26 | 0.708 | 0.26 |
| | Metric B | 19.51 | 0.667 | 0.39 |
| | Metric C | 16.13 | 0.833 | 0.06 |
| | Metric D | 19.23 | 0.694 | 0.29 |
| | Metric E | 19.63 | 0.639 | 0.47 |
| | Metric F | 16.68 | 0.778 | 0.13 |
| | Metric G | 20.26 | 0.611 | 0.59 |
| 2 | Metric C + A | 16.31 | 0.861 | 0.041 |
| | Metric C + B | 15.86 | 0.917 | 0.015 |
| | Metric C + D | 11.60 | 0.861 | 0.045 |
| | Metric C + E | 14.42 | 0.972 | 0.004 |
| | Metric C + F | 13.99 | 0.972 | 0.004 |
| | Metric C + G | 17.19 | 1.000 | 0.002 |
| | Metric F + A | 18.58 | 0.806 | 0.093 |
| | Metric F + B | 18.56 | 0.806 | 0.093 |
| | Metric F + C | 13.99 | 0.972 | 0.004 |
| | Metric F + D | 18.45 | 0.806 | 0.093 |
| | Metric F + E | 18.41 | 0.861 | 0.041 |
| | Metric F + G | 16.09 | 0.889 | 0.026 |
| 3 | **Metric D + C + A** | **14.67** | **1.000** | **0.002** |
| | Metric D + C + B | 15.24 | 1.000 | 0.002 |
| | Metric D + C + E | 16.12 | 0.944 | 0.009 |
| | Metric D + C + F | 15.56 | 0.972 | 0.004 |
| | Metric D + C + G | 19.66 | 1.00 | 0.002 |
| | Metric C + F + A | 15.05 | 0.972 | 0.004 |
| | Metric C + F + B | 14.71 | 0.972 | 0.004 |
| | Metric C + F + D | 15.56 | 0.972 | 0.004 |
| | Metric C + F + E | 13.55 | 0.972 | 0.004 |
| | Metric C + F + G | 15.84 | 0.944 | 0.009 |
| 4 | Metric D + C + A + B | 16.06 | 1.000 | 0.002 |
| | Metric D + C + A + E | 16.03 | 1.000 | 0.002 |
| | Metric D + C + A + F | 14.80 | 1.000 | 0.002 |
| | Metric D + C + A + G | 15.57 | 1.000 | 0.002 |
| | Metric C + F + E + A | 15.57 | 1.000 | 0.002 |
| | Metric C + F + E + B | 15.57 | 1.000 | 0.002 |
| | Metric C + F + E + D | 15.57 | 1.000 | 0.002 |
| | Metric C + F + E + G | 15.57 | 1.000 | 0.002 |
| 5 | Metric D + C + A + F+ B | 20.05 | 0.944 | 0.009 |
| | Metric D + C + A + F+ E | 19.02 | 0.972 | 0.004 |
| | Metric D + C + A + F+ G | 17.44 | 0.944 | 0.009 |
| 7 | Metric A + B + C + D + E + F + G | 21.21 | 1.000 | 0.002 |

**Fig. S4. Logistic regression model of reduced mAb concentrations.** Tables show the results of the logistic regression analysis for corresponding ROC plots in Figure 3A.

**A** *Signal metrics analysis*

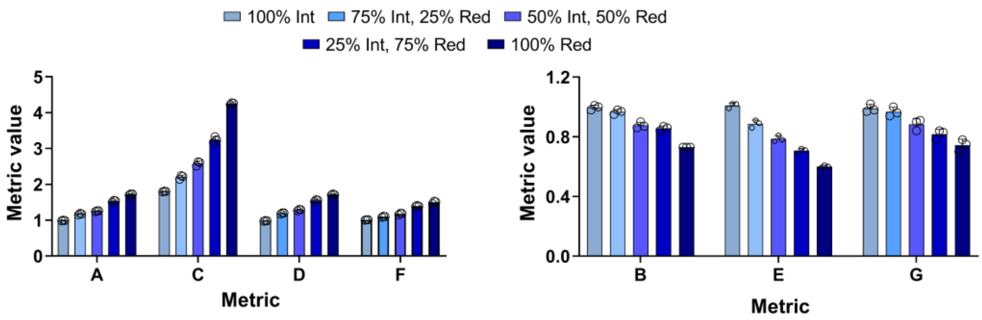


| Metric | 0% Reduced | 25% Reduced | 50% Reduced | 75% Reduced | 100% Reduced |
|---|---|---|---|---|---|
| A | 0.99 ± 0.01 | 1.17 ± 0.02 | 1.25 ± 0.01 | 1.53 ± 0.02 | 1.72 ± 0.02 |
| B | 1.00 ± 0.02 | 0.96 ± 0.02 | 0.88 ± 0.02 | 0.85 ± 0.02 | 0.73 ± 0.00 |
| C | 1.80 ± 0.01 | 2.20 ± 0.05 | 2.58 ± 0.06 | 3.24 ± 0.08 | 4.25 ± 0.03 |
| D | 0.98 ± 0.01 | 1.19 ± 0.02 | 1.28 ± 0.02 | 1.55 ± 0.03 | 1.71 ± 0.01 |
| E | 1.01 ± 0.02 | 0.89 ± 0.02 | 0.78 ± 0.02 | 0.71 ± 0.01 | 0.60 ± 0.01 |
| F | 1.01 ± 0.01 | 1.10 ± 0.02 | 1.18 ± 0.03 | 1.39 ± 0.02 | 1.50 ± 0.04 |
| G | 0.99 ± 0.02 | 0.97 ± 0.03 | 0.88 ± 0.04 | 0.82 ± 0.03 | 0.74 ± 0.04 |

**B** *Linear regression*

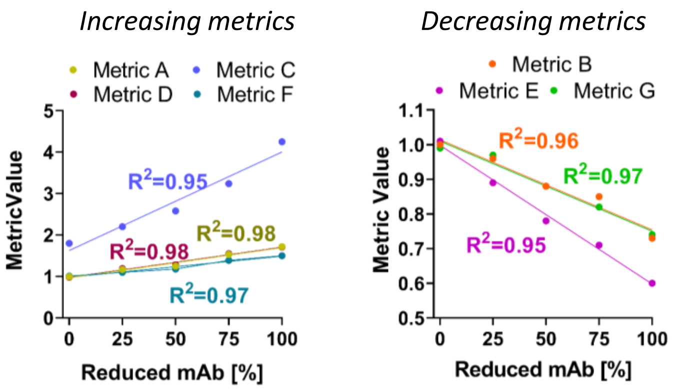


**Fig. S5. Varying reduced mAb composition. (A)** Bar graphs and tables of signal metrics for corresponding data in Figure 3B. Error bars in plot represent the standard deviation. **(B)** Linear regression of CV and CC signal metrics across all reduced mAb compositions evaluated.

|  | Buffer |  | Fresh media |  | Conditioned media |  |
|---|---|---|---|---|---|---|
| Metric | Intact | Reduced | Intact | Reduced | Intact | Reduced |
| A | 1.05 ± 0.05 | 1.45 ± 0.05 | 1.03 ± 0.06 | 1.39 ± 0.06 | 1.00 ± 0.03 | 1.16 ± 0.03 |
| B | 1.05 ± 0.01 | 0.61 ± 0.01 | 1.03 ± 0.11 | 0.49 ± 0.11 | 1.00 ± 0.03 | 0.59 ± 0.03 |
| D | 1.02 ± 0.06 | 1.65 ± 0.06 | 1.00 ± 0.04 | 1.56 ± 0.04 | 0.98 ± 0.05 | 1.34 ± 0.05 |
| E | 1.06 ± 0.02 | 0.57 ± 0.02 | 1.04 ± 0.10 | 0.44 ± 0.10 | 1.01 ± 0.07 | 0.54 ± 0.07 |
| F | 1.03 ± 0.02 | 2.36 ± 0.03 | 1.01 ± 0.01 | 2.38 ± 0.08 | 1.05 ± 0.02 | 1.88 ± 0.05 |
| G | 1.05 ± 0.01 | 0.93 ± 0.03 | 0.98 ± 0.01 | 0.68 ± 0.04 | 1.07 ± 0.02 | 0.79 ± 0.03 |

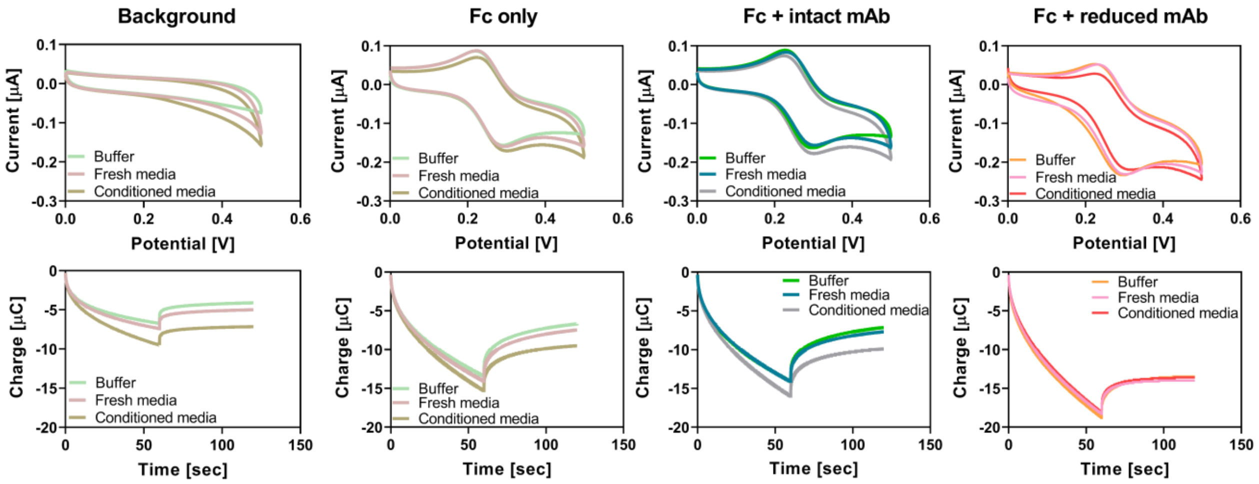


**Fig. S6. Varying background composition of reduced mAb.** Table of values for signal metrics ($n$ = 3). Error represents the standard deviation. Plots show comparison of background (buffer only), Fc only, Fc + intact mAb, and Fc + reduced mAb diluted in buffer, fresh media, or conditioned media.

**A** *Antibody oxidation*

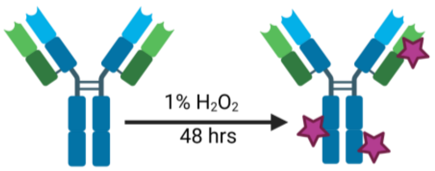


**B** *NISTmAb structure*

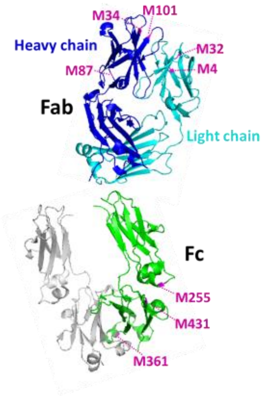


**C** *NISTmAb peptide mapping*

| | | Met Oxidation [%] | |
|---|---|---|---|
| **Chain** | **Reside** | **Control** | **Oxidized** |
| Heavy | M34 | 7.79 | 99.48 |
| Heavy | M87 | 1.28 | 2.71 |
| Heavy | M101 | 7.09 | 99.20 |
| Heavy | M255 | 7.59 | 99.90 |
| Heavy | M361 | 0.16 | 95.35 |
| Heavy | M431 | 2.16 | 99.32 |
| Light | M4 | 9.12 | 98.41 |
| Light | M32 | 2.60 | 6.31 |
| **Total % Met Oxidation** | | **5.52** | **84.07** |

**D** *SASA analysis*

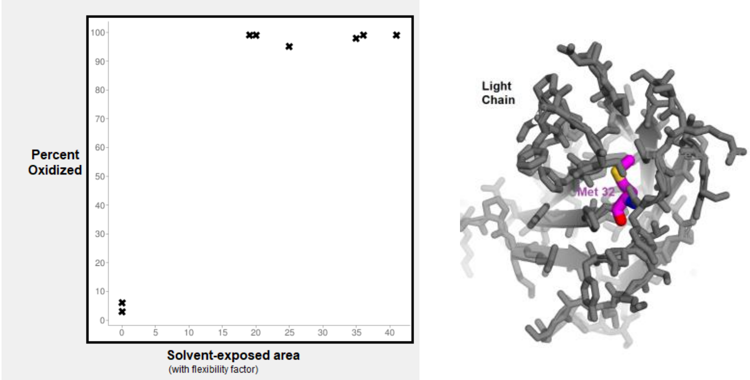


**E** *Durvalumab peptide mapping*

| | | Met Oxidation [%] | |
|---|---|---|---|
| **Chain** | **Reside** | **Control** | **Oxidized** |
| Heavy | M34 | 2.18 | 3.48 |
| Heavy | M83 | 0.78 | 1.49 |
| Heavy | M256 | 5.38 | 99.89 |
| Heavy | M362 | 1.86 | 98.23 |
| Heavy | M432 | 2.81 | 99.14 |
| **Total % Met Oxidation** | | **2.52** | **54.76** |

**Fig. S7. Methods to create and analyze oxidized mAb variants. (A)** Method for antibody oxidation. **(B)** Eight methionine residues of NISTmAb (one heavy-light chain pair) are highlighted in pink (PDB: 5K8A; the Fab domain and 5VGP; the Fc domain). **(C)** LC-MS/MS peptide mapping results show methionine residues of control (non-oxidized) NISTmAb (one heavy-light chain pair) are not significantly oxidized. For NISTmAb that has undergone the oxidation protocol, six methionine residues are completely oxidized (> 98%). **(D)** Solvent-Accessible Surface Area (SASA) results indicate that the solvent accessible area for methionine residues that did not become oxidized was lower than residues that became oxidized. The correlation between the SASA and the measured oxidation values in the plot was 0.87. Graphic shows that non-oxidized methionine residues (e.g., M32) are buried in the structure (light chain of the Fab) and thus shielded from oxidation by hydrogen peroxide. **(E)** LC-MS/MS peptide mapping confirms that durvalumab (one heavy-light chain pair) has three methionine residues that are oxidized.

**A** *Control mAb comparison with Ir*

**B** *Signal metrics*

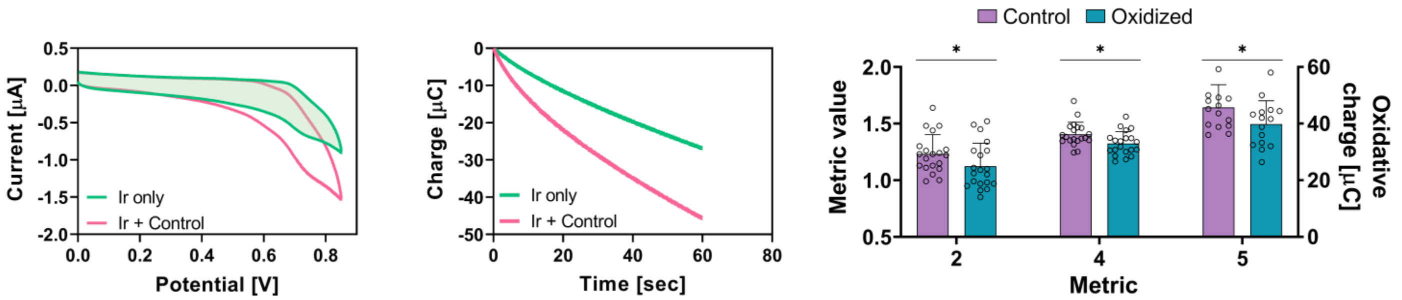


**C** *Logistic regression model*

| # of metrics | Combination | AIC | AUC | *p*-value |
|---|---|---|---|---|
| 1 | Metric 1 | 43.17 | 0.644 | 0.180 |
| | Metric 2 | 43.41 | 0.658 | 0.150 |
| | Metric 3 | 41.58 | 0.707 | 0.056 |
| | Metric 4 | 41.51 | 0.702 | 0.061 |
| | Metric 5 | 41.64 | 0.700 | 0.065 |
| | Metric 6 | 37.16 | 0.813 | 0.004 |
| 2 | Metric 6 + 1 | 38.82 | 0.84 | 0.001 |
| | Metric 6 + 2 | 39.16 | 0.818 | 0.002 |
| | Metric 6 + 3 | 36.18 | 0.840 | 0.001 |
| | Metric 6 + 4 | 36.50 | 0.831 | 0.001 |
| | Metric 6 + 5 | 39.06 | 0.800 | 0.004 |
| 3 | **Metric 6 + 3 + 1** | **32.14** | **0.902** | **0.00005** |
| | Metric 6 + 3 + 2 | 32.87 | 0.876 | 0.002 |
| | Metric 6 + 3 + 4 | 38.15 | 0.844 | 0.008 |
| | Metric 6 + 3 + 5 | 37.49 | 0.831 | 0.001 |
| 4 | Metric 6 + 3 + 1 + 2 | 34.13 | 0.902 | 0.00005 |
| | Metric 6 + 3 + 1 + 4 | 34.08 | 0.902 | 0.00005 |
| | Metric 6 + 3 + 1 + 5 | 33.83 | 0.889 | 0.0001 |
| 6 | Metric 1 + 2 + 3 + 4 + 5 + 6 | 37.30 | 0.907 | 0.00004 |

**Fig. S8. MEP of oxidized NISTmAb. (A)** Control NISTmAb (0.25 g/L) comparison with Ir (50 µmol/L) only. **(B)** Bar graph indicates results of signal metrics analysis. Error bars represent the standard deviation. **(C)** Table of logistical regression analysis.

*Varying mAb concentration*

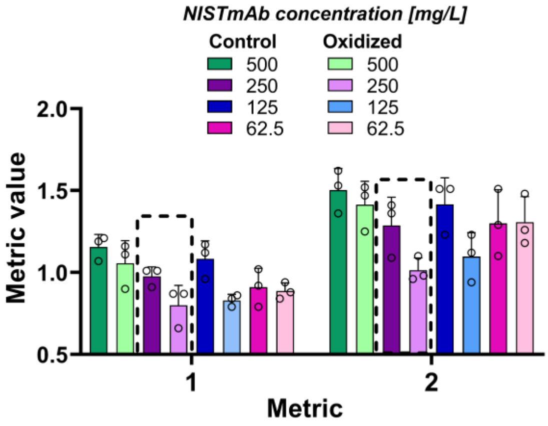


**Metric 1**

| mAb [g/L] | Intact | Oxidized | Difference |
|---|---|---|---|
| 500 | 1.16 ± 0.07 | 1.06 ± 0.14 | - 0.10 |
| 250 | 0.98 ± 0.06 | 0.80 ± 0.12 | - 0.18 |
| 125 | 1.08 ± 0.11 | 0.83 ± 0.04 | - 0.25 |
| 62.5 | 0.91 ± 0.11 | 0.89 ± 0.05 | - 0.02 |

**Metric 2**

| mAb [g/L] | Intact | Oxidized | Difference |
|---|---|---|---|
| 500 | 1.50 ± 0.13 | 1.41 ± 0.15 | - 0.09 |
| 250 | 1.29 ± 0.17 | 1.01 ± 0.07 | - 0.28 |
| 125 | 1.42 ± 0.16 | 1.10 ± 0.15 | - 0.32 |
| 62.5 | 1.30 ± 0.21 | 1.31 ± 0.16 | + 0.01 |

*Varying Ir concentration*

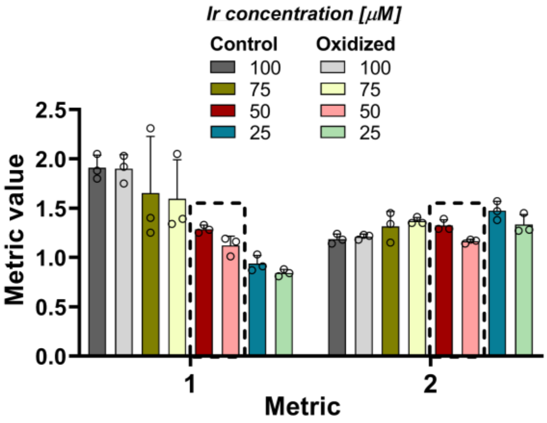


**Metric 1**

| Ir [μM] | Intact | Oxidized | Difference |
|---|---|---|---|
| 100 | 1.91 ± 0.08 | 1.90 ± 0.14 | - 0.01 |
| 75 | 1.65 ± 0.04 | 1.59 ± 0.39 | - 0.06 |
| 50 | 1.29 ± 0.57 | 1.12 ± 0.10 | - 0.17 |
| 25 | 0.94 ± 0.13 | 0.84 ± 0.03 | - 0.10 |

**Metric 2**

| Ir [μM] | Intact | Oxidized | Difference |
|---|---|---|---|
| 100 | 1.19 ± 0.05 | 1.21 ± 0.03 | + 0.02 |
| 75 | 1.31 ± 0.15 | 1.37 ± 0.04 | + 0.06 |
| 50 | 1.32 ± 0.06 | 1.16 ± 0.02 | - 0.16 |
| 25 | 1.48 ± 0.09 | 1.33 ± 0.10 | - 0.15 |

**Fig. S9. Optimization of analysis.** Varied mAb concentration (with 50 µmol/L Ir) and Ir concentration (with 0.25 g/L NISTmAb) to demonstrate effort to obtain an “optimal” ratio of mAb:Ir is needed to discern between control and oxidized mAbs ($n$ = 3). Error bars in the plots represent the standard deviation.

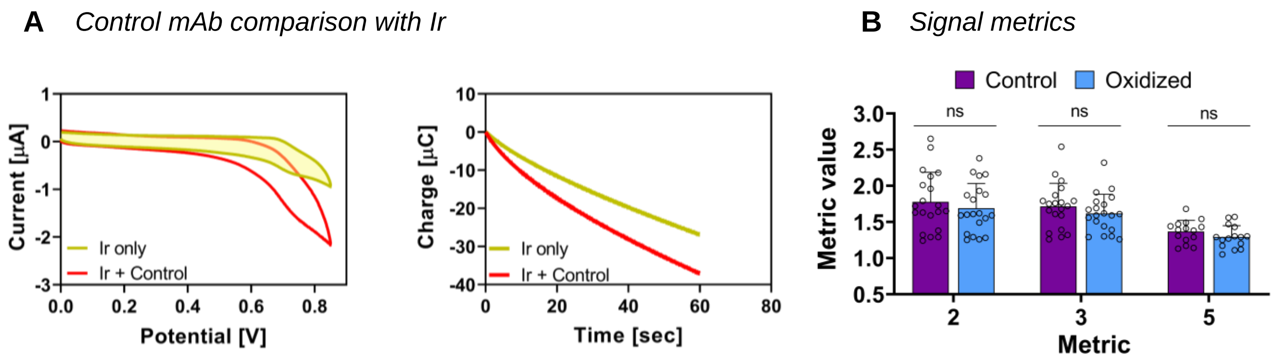


**C** *Logistic regression model*

| # of metrics | Combination | AIC | AUC | *p*-value |
|---|---|---|---|---|
| 1 | Metric 1 | 44.02 | 0.636 | 0.220 |
| | Metric 2 | 44.61 | 0.631 | 0.230 |
| | Metric 3 | 43.29 | 0.687 | 0.085 |
| | Metric 4 | 44.04 | 0.642 | 0.190 |
| | Metric 5 | 43.75 | 0.640 | 0.200 |
| | Metric 6 | 38.89 | 0.771 | 0.012 |
| 2 | **Metric 6 + 1** | **39.43** | **0.796** | **0.005** |
| | Metric 6 + 2 | 39.77 | 0.778 | 0.008 |
| | Metric 6 + 3 | 39.88 | 0.787 | 0.006 |
| | Metric 6 + 4 | 40.45 | 0.764 | 0.013 |
| | Metric 6 + 5 | 40.86 | 0.756 | 0.016 |
| 3 | Metric 6 + 1 + 2 | 41.40 | 0.791 | 0.005 |
| | Metric 6 + 1 + 3 | 41.42 | 0.796 | 0.004 |
| | Metric 6 + 1 + 4 | 40.90 | 0.791 | 0.005 |
| | Metric 6 + 1 + 5 | 40.98 | 0.800 | 0.004 |
| 4 | Metric 6 + 1 + 4 + 2 | 42.70 | 0.796 | 0.004 |
| 6 | Metric 1 + 2 + 3 + 4 + 5 + 6 | 44.07 | 0.836 | 0.001 |

**Fig. S10. MEP of oxidized durvalumab.** (A) Control durvalumab (0.25 g/L) comparison with Ir (50 µmol/L). **(B)** Bar graph indicates results of signal metrics analysis. Error bars in plot represent the standard deviation. **(C)** Table of logistical regression analysis.
